# Specific interaction of Penetratin with cell surface partners measured with biomembrane force probe

**DOI:** 10.1101/2020.07.21.215111

**Authors:** P Soule, F Illien, S Kulifaj, A Joliot, C Gourier, S Sagan, S Cribier, N Rodriguez

## Abstract

Penetratin is a Cell Penetrating Peptide able to cross the cell plasma membrane possibly bound to a cargo molecule to be delivered into the cell. The mechanism of its entry is poorly known. A key to a molecular description of this mechanism is to identify the partners of Penetratin at the cell surface during its adhesion and internalization. We used the Biomembrane Force Probe to identify the partners during the first second of adhesion of Penetratin on the cell plasma membrane. We evidenced that heparan sulfates are the first partners after contact as well as unknown partners hidden by sialic acids. Experiments of binding of Penetratin on vesicles bearing charged or sulfated lipids showed no adhesion pointing that a negatively charged partner is not enough and there is a specificity for certain chemical groups bearing the charges. A model of the measured forces of interaction enabled to determine the adhesion energy of a Penetratin with heparan sulfates on a cell to be in the range 18 to 22 *k*_*B*_*T*.

## INTRODUCTION

### CPP and its potential partners

Cell Penetrating Peptides (CPPs) are a family of short peptides, highly enriched in basic residues, sharing the ability to enter into cells. This ability can be retained when bound to cargos and they are thus potentially very promising to deliver exogenous molecules into cells (Kurrikoff et al., 2015, 2016). They have been shown to enter cells by two pathways: endocytosis and direct translocation through the plasma membrane (Bechara and Sagan, 2013; Jiao et al., 2009). However the molecular mechanism of this entry is poorly understood. A major question regarding this mechanism is the nature of the protein, lipid or polysaccharide partners of the CPPs at each step of the interaction of the CPPs with the plasma membrane: adhesion at the membrane, membrane deformation and membrane crossing. This study focused on a CPP, the Penetratin, corresponding to the third helix of the Antennapedia homeodomain (Derossi et al., 1994). Penetratin is among the first discovered CPPs and the subject of many studies to understand the mechanism of internalization of CPPs (Dupont et al., 2015). The aim was to determine the partners and quantify the energy of adhesion of the Penetratin at the adhesion step of the Penetratin on the plasma membrane of CHO cells. Several negatively charged molecules are good candidates to bind the positively charged Penetratin: (i) surface proteins especially if negatively charged (ii) negatively charged lipids and sulfate/phosphate/carboxylate bearing lipids (this latter chemical groups can form bonds with guadininium groups on arginine residues from CPPs; the Penetratin bears 3 arginine residues) (Rothbard et al., 2004) (iii) polysaccharides worn by glycosaminoglycans (GAGs) especially when sulfated such as chondroitin sulfates or heparane sulfates (Console et al., 2003).

### Techniques to identify partners

Several techniques have already been used to identify CPP partners. Some enable a determination of the energy of the CPP-partner adhesion; others use cells containing the partners instead of isolated partners or reconstituted membranes. We will show that the Biomembrane Force Probe technique used in this study combined both advantages.

Quantification by mass spectrometry of the CPPs bound to the membrane and internalized into cells (Burlina et al., 2005, 2006) and the comparison between wild type CHO cells and CHO cells lacking surface GAGs demonstrated that these GAGs did not impact the amount of Penetratin bound on the cell membrane but had a strong effect on its internalization (mainly due to endocytosis for wild type cells and direct translocation for GAG deficient cells) (Jiao et al., 2009). These results have been confirmed recently by another method of quantification using fluorimetry (Illien et al., 2016).

The previous methods enable to detect Penetratin adhesion but do not quantify the energy of interaction. This latter parameter can be obtained with isothermal calorimetry (ITC): the constant of dissociation of Penetratin with several sugars -heparin and several chondroitin sulfates-are in the tens-hundreds of nanomolars range with no noticeable preference for heparin (Bechara et al., 2013; Ziegler and Seelig, 2008).

Another technique combines quantitative determination of dissociation constant between a CPP and its partners and the presence of a reconstituted membrane as a target of the CPP: Plasmon Surface Waveguide Resonance. It enabled to determine a dissociation constant of Penetratin on membrane reconstituted from different cell lines. The 10 nM dissociation constant of Penetratin obtained for membrane originating from WT CHO cells was significantly lower than that from GAG deficient or sialic acids (SAs) deficient cells (Alves et al., 2011).

The technique we used in this study allows both to seek for the partners of the Penetratin on intact cells and to obtain quantitative data about the energetics of the adhesion. It relied on the use of the Biomembrane Force Probe (BFP) (Evans et al., 2004). BFP enabled to detect the force exerted between few Penetratins and a target cell or vesicle after a short contact leading to an adhesion or not depending on the presence of different partners at the cell/vesicle surface. It is used here to find the main partners of Penetratin during the first second of its adhesion with the membrane of a target CHO cell and to quantify the energy of this adhesion.

## MATERIALS AND METHODS

### Peptide

Biotin-Aminopentanoyl-Penetratin was obtained in our laboratory by solid-phase synthesis using the Boc strategy. Penetratin was purified by RP-HPLC and further checked by mass spectrometry.

### Cell culture

Wild type Chinese Hamster Ovary CHO-K1 cells, xylose-transferase- or GAG-deficient CHO-pgsA745 cells and sialic acid deficient CHO-lec2 cells (ATCC, LGC Standards S.a.r.l. - France) were cultured in F12 growth medium (DMEM-F12) supplemented with 10% fetal calf serum (FCS), penicillin (100,000 IU/L), streptomycin (100,000 IU/L), and amphotericin B (1 mg/L) in a humidified atmosphere containing 5% CO_2_ at 37 °C.

### Cell detachment

CHO cells are adherent cells which need to be detached before the BFP experiment. The cells are detached by 0.5mM EDTA incubation in PBS for 5 min at 37°C. After detachment the cells are spun down to the right concentration in PBS.

### Heparinase III treatment

After detachment the cells are spun down to the right concentration in PBS and incubated for one hour at 37°C with 40nU per cell (1x treatment) or 200nU per cell (5x treatment) of heparinase III from Flavobacterium heparinum (Sigma Aldrich). The treated cells are then immediately used for a < 2h BFP experiment.

### Giant vesicles

20µL of a 20µg/µL lipid solution in chloroform are deposited on the bottom of a rough teflon dish. The droplet is spread and the chloroform evaporates under vacuum for 30 min. The dish is then filled with 300 mM sucrose in water solution and the lipid film swells overnight at 37°C. In this condition vesicles with diameter ∼20µm are obtained. Given the condition of their formation, they are expected to be multilamellar.

### Biomembrane Force Probe

The principle of our Biomembrane Force Probe experiment is to bring into contact a bead bearing Penetratins with a target cell or vesicle and to measure the force needed to separate the bead from the target when pulling the bead away from the cell. If a non zero force is recorded it is due to the adhesion of few Penetratins with partners on the target membrane. To measure this force the deformation of a red blood cell acting as a calibrated spring is monitored. The red blood cell is held by a glass micropipette whose position is controlled by a piezoelectric translator (Physik Instruments) and bears the bead on which Penetratins are grafted (figure 1A). To determine the deformation of the red blood cell, the positions of its extremities need to be determined. The relative position of its base is controlled by the piezoelectric device with a nanometric accuracy. The position of its other side is deduced from the position of the adherent bead which is tracked by videomicroscopy with an accuracy ∼20nm: the images are acquired at 300Hz by a u-eye camera (Stemmer Imaging) mounted on a microscope and analyzed in real time by an home-made software. The force exerted on the bead is deduced from the deformation of the red blood cell through the spring stiffness inferred from geometric parameters of the red blood cell-bead complex and the controlled aspiration of the red blood cell through the micropipette (Evans et al., 1995).

**Figure 1:**
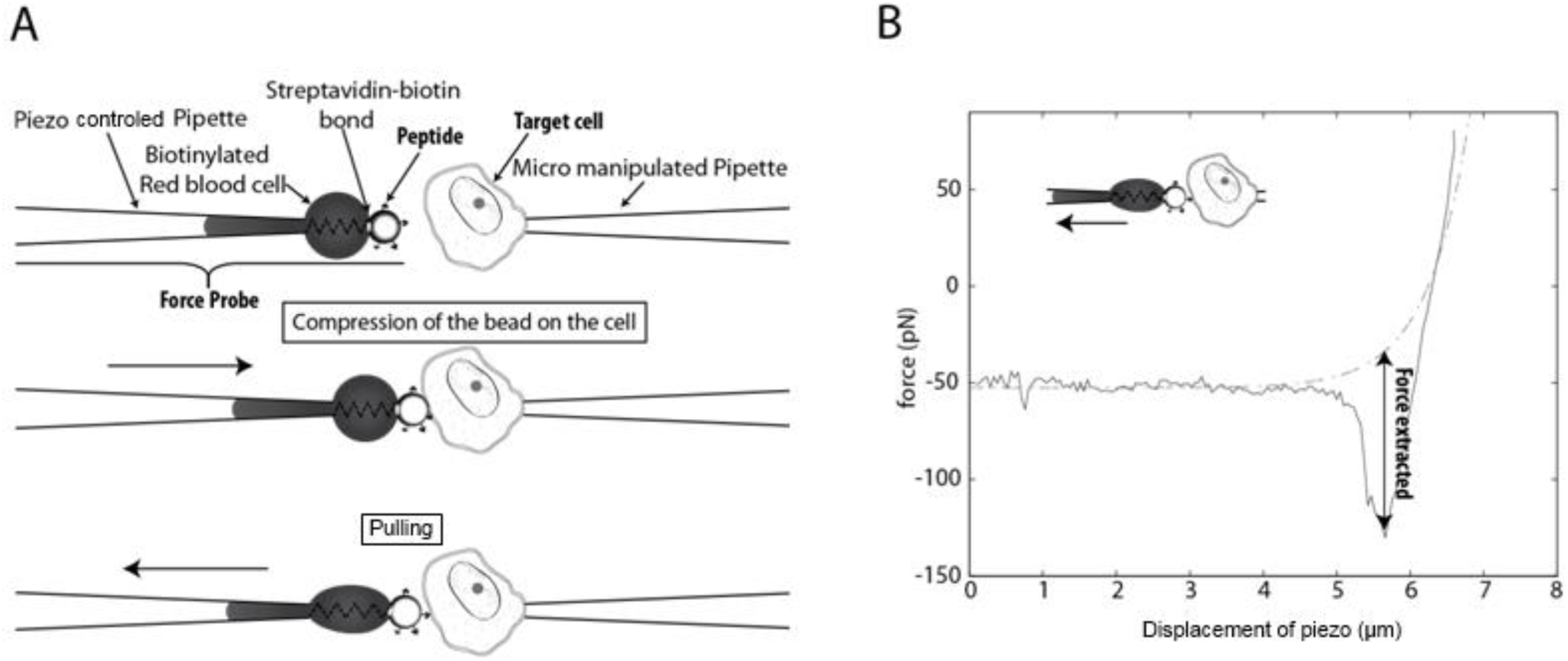
Principle of Biomembrane Force Probe. A: Scheme of the Biomembrane Force probe. Top: elements of the BFP; Middle: pushing phase, the bead is pressed on the target cell; Bottom: pulling phase, an adhesion occurred, the red blood cell is stretched before rupture. B: Example of force curve measured during the pulling phase. An adhesion is observed. The adhesion force (depth of the well) is measured by subtraction from the measured force curve of a fit of the beginning and end of the force curve.

During an experiment series of way-back sequences are imposed to the piezotranslator. (i) The bead is pushed into contact with the still target cell or vesicle (held by another micropipette facing the other one) until an imposed 80pN force is reached. (ii) The contact is maintained 400 ms. (iii) The bead is pulled away from the cell/vesicle; the unloading rate during the withdrawal of the bead is set at 4800pN/s. If an adhesion occurred between the molecules on the bead and the target membrane, a rupture force is necessary to detach the bead from the cell/vesicle. Figure 1B presents an example of force curve during the pulling phase. On this example a rupture force is detected at the beginning of the pulling.

For each condition (type of cell or vesicle/Penetratin or not on the bead/possible treatment of the cell) ∼1000 to 3000 curves were obtained with ∼5 to 15 different cells/vesicles.

### Red blood cell biotinylation

The red blood cells are rinsed twice with PBS buffer and twice with bicarbonate buffer (carbonate-bicarbonate 100 mM, pH 8.5) before an incubation with 2mg/mL Biotin-30 kDa PEG-Succinimidyl ester (Interchim) in bicarbonate buffer for 1h. They are then rinsed three times in Tris buffer (Tris 25 mM, NaCl 150 mM, pH 7.5) and incubated for another 30min in Tris buffer before being rinsed three times with PBS buffer. The red blood cells are stored at 4°C and used within few weeks.

### Beads preparation

The glass beads are 3µm diameter and coated with streptavidin (Jégou et al., 2008). To bind Biotin-Aminopentanoyl-Penetratin we incubated 10^6^ beads in 1mL water with 20µM Penetratin for 5min and the beads are washed several times with water to get rid of the excess of peptide.

### Pipette formation

Micropipettes are obtained by elongating borosilicate glass capillaries (1-mm outer diameter, 0.78-mm inner diameter, Harvard Apparatus, Holliston, MA) with a micropipette puller (P-2000, Sutter Instrument, Novato, CA). The pipettes holding the red blood cell are then forged to obtain a smooth open tip with an inner diameter of ∼2µm. The inner diameter of the tip of the pipettes holding the cell or the vesicle is ∼2.5µm.

### Data analysis

The data were analyzed using a homemade algorithm. The algorithm is programmed on MATLAB (The MathWorks) and consists into a fit of the force curve during the pulling using an exponential plus a linear term apart from the position of a possible well stemming from a possible adhesion force. If no adhesion is detected this fit follows closely the pulling force curve. If an adhesion occurred, there is a well at the beginning of the pulling force curve due to the rupture force needed to detach the adherent bead from the cell/vesicle. The depth of the well is calculated by subtraction of the fit to the force curve. The noise is estimated from the planar half of the force curve. If the depth of the well is above 3 times the noise and above 30pN, an adhesion is considered to have been detected and the depth of the well is named “adhesion force” (figure 1B).

### Determination of the adhesion energies

The aim of the model presented below is to compute the distribution of rupture forces for one or two Penetratins bound to their partners on the target cell. This model relies on the hypothesis of the interaction energy between Penetratin and its partner to be a single well of depth *E*_*a*_ and width *x*_*w*_. To obtain the distribution of rupture force predicted by the model, a probability of rupture per unit of time is computed and compared with random draws to determine if the bond is ruptured during a time interval. This process is repeated until the rupture of the bond and the force at rupture is then registered. The force at time t (possibly divided between two bonds when two Penetratins are bound) is *f=αt* where *α* = 4800*pN /s* is the load rate. The energy of adhesion per Penetratin *E*_*a*_ is drawn from a normal distribution centered on *E*_*am*_ and of standard deviation *σ*_*a*_.

We distinguish two cases of probabilities p1 and p2 respectively: Case 1: One penetratin is bound:

The probability *dP* of rupture of the bond at time t during dt is given by:

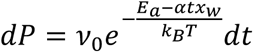

*v*_0_ is a frequency of order of magnitude 10^10^ *s*^−1^ (Merkel et al., 1999; Pincet and Husson, 2005).

Case 2: Two Penetratins are initially bound:

The probability *dP*of rupture of one of the two bonds at time t during dt is given by:

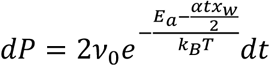

After rupture of the first bond the probability *dP* of rupture of the last bond at time t during dt is given by:

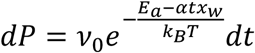

The time of rupture of the unique or last bond is simulated and the corresponding rupture forces is deduced from *f* = *αt*. This process is repeated and the distribution of rupture forces is deduced from (p1+ p2)*10^6^ trials (p1*10^6^ trials with one Penetratin initially bound and p2*10^6^ trials with two Penetratins initially bound).

## RESULTS

### Biomembrane Force Probe allows the detection of the adhesion between few penetratins and some partners at the membrane of a CHO cell

The Biomembrane Force Probe allows the measurement of the frequency of the adhesions between a bead coated with Penetratin peptides and a target cell as well as the force necessary to separate the bound peptide(s) from the membrane. In the proper conditions an adhesion of the bead on the target cells is not observed for all bead-cell contacts. To assess if this technique is suitable to study the interaction between Penetratin and the CHO cell membrane, it is necessary to tune the Penetratin bead coverage so that the frequency of adhesion is significantly different from the frequency of unspecific adhesions (adhesion with an uncovered bead). In the absence of Penetratin on the bead there were some unspecific adhesions between the bead and the CHO cell membrane: it occurred for 13.6% of the contacts (figure 2A). With the proper coverage of the bead by Penetratins, the frequency of adhesion between a Penetratin coated bead and a CHO cell rose to 38.1% (significantly different from 13.6%, p-value<10^−5^). In 24.5% of the contacts between a coated bead and a cell the interaction can then be attributed to the formation of one or several bonds between one or several Penetratin peptides and the membrane (this specific rate is obtained by subtraction of the rates with and without Penetratin on the bead). The Biomembrane Force Probe is therefore a suitable technique to study the interactions of Penetratin with a cell membrane. In the following experiments specific interactions are defined as the total interaction in the presence of Penetratin minus the interactions with non coated beads.

**Figure 2:**
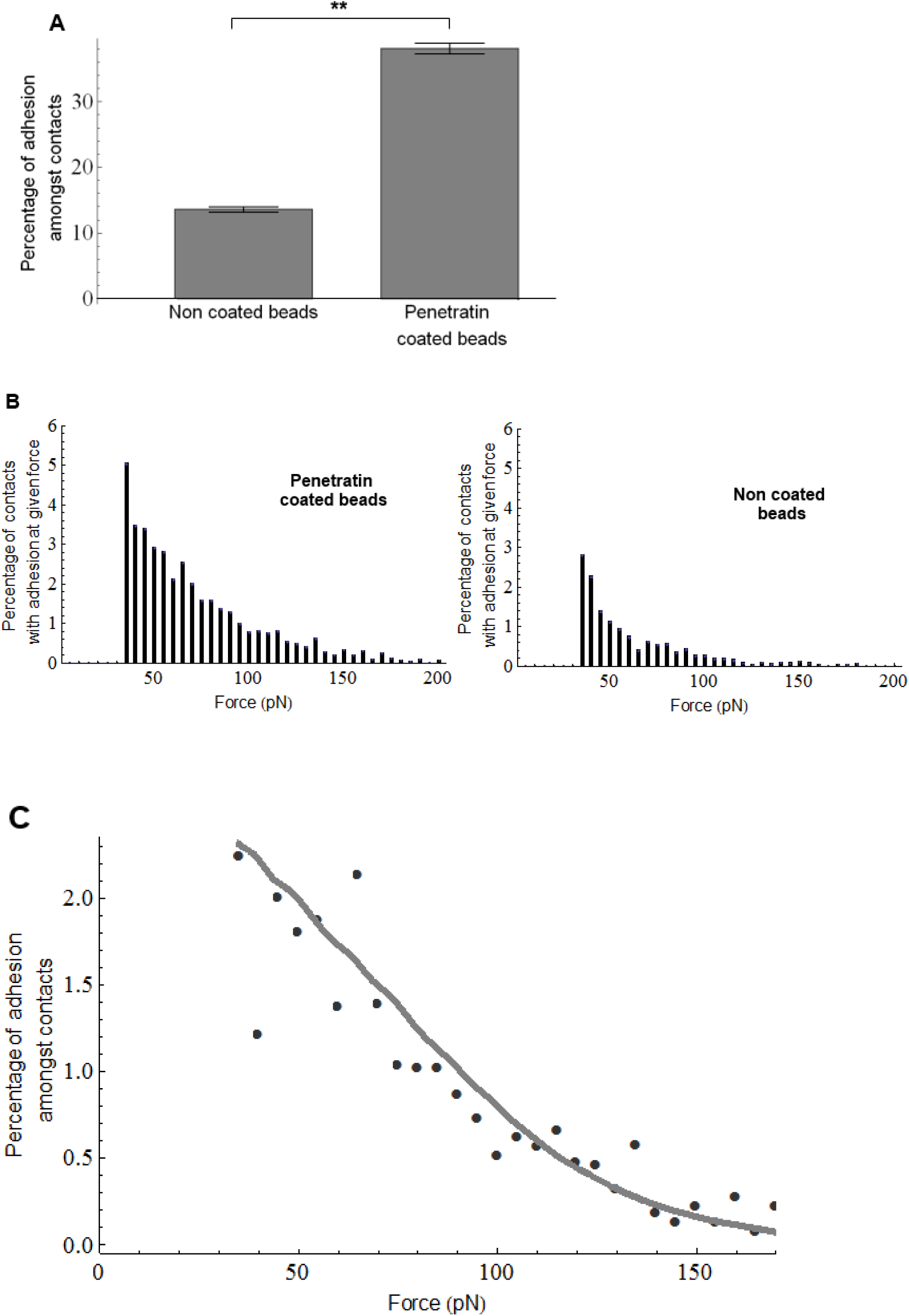
Specific Penetratin adhesion with CHO wild type cells is detected. A: Penetratin coated beads display a significantly higher adhesion rate than non coated beads. B: Force distribution curve of Penetratin coated beads and non coated beads. C: Comparison between the measured specific to Penetratin on WT CHO cell force distribution (black dots) and a simulated simple well model (gray line). The parameters of the best fit model are : *E*_*am*_ = 19.9 *k*_*B*_*T, σ*_*a*_ = 1.9 *k*_*B*_*T, v*_0_ = 10^10^ *s*^−1^, *p*1 = 0.33, *p*2 = 0.1 and *x*_*w*_ = 2Å.

The study of the force distribution of the specific adhesions (obtained by subtraction of the unspecific force distribution to the Penetratin covered bead force distribution (figure 2B)) can be used to infer the energy of adhesion of Penetratin with its partners at the surface of CHO cell. The determination of this parameter from BFP data is not straightforward but an estimation can be attempted. For this purpose, we use a simple energetic landscape model of the bond between Penetratin and its partners at the membrane: a simple well with an activation energy barrier of magnitude *E*_*a*_ and width *x*_*w*_. Because for most of the contacts we detect no specific adhesion, only 0, 1 or two Penetratins are significantly likely to bind for each contact with probability p0, p1 and p2 respectively. The force applied during the pulling of the bond is *f=αt* where *α* is the load rate. The height of the energy barrier at time t is then *E*_*a*_*-αtx*_*w*_when one Penetratin is bound and 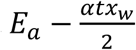 when two Penetratins are bound. To take into account the fact that Penetratin may bind not always with the same orientation on its partners, the values of *E*_*a*_ obey a distribution of values rather than a single value. We propose as a simple model a normal distribution centered on *E*_*am*_ and of standard deviation *σ*_*a*_. Taking all these parameters into account a probability of rupture of a bond per unit of time can be computed (supplementary material) and the distribution of rupture forces can be simulated as described in the method section and compared with the experimental distribution of force (figure 2C). We found the adhesion energy of the Penetratin with its partner to be *E*_*a*_*=*18 *to* 22*k*_*B*_*T*.

### The heparan sulfates on the cell membrane are responsible for the specific interaction of penetratins with the cell membrane

We detected a specific partner of Penetratin on the CHO cell surface and determined its adhesion energy, we then turned to the determination of this partner.

The GAGs are known to be partners of Penetratin when it binds cell membranes (Console et al., 2003). We have tested the interaction of beads bearing Penetratins with pgsA745 CHO cells (GAG deficient cells), which are cells lacking most of their GAGs.

The frequency of specific interactions of Penetratin coated beads with GAG deficient CHO cells is not significantly different from the frequency of specific interaction of these beads with wild type CHO cells (figure 3A: 24.5% and 24.7% respectively). However GAG deficient CHO cells lack most of their GAGs but not all because the xylose transferase activity in these cells is strongly reduced but not abolished (Esko et al., 1985). The residual GAGs may be dense enough to find Penetratins on the bead in a BFP experiment. We therefore treated our GAG deficient CHO cells with heparinase III, an enzyme which specifically degraded heparan sulfates (HSs), which are GAGs that are known to interact with cell penetrating peptides (Bechara et al., 2013). On GAG deficient cells, the frequency of specific interactions was completely abolished (0% of specific interaction for heparinase III treated GAG deficient CHO cells). We can deduce that the specific interactions measured between GAG deficient CHO cells and Penetratin coated beads were due to the residual HSs at the surface of the cells.

**Figure 3:**
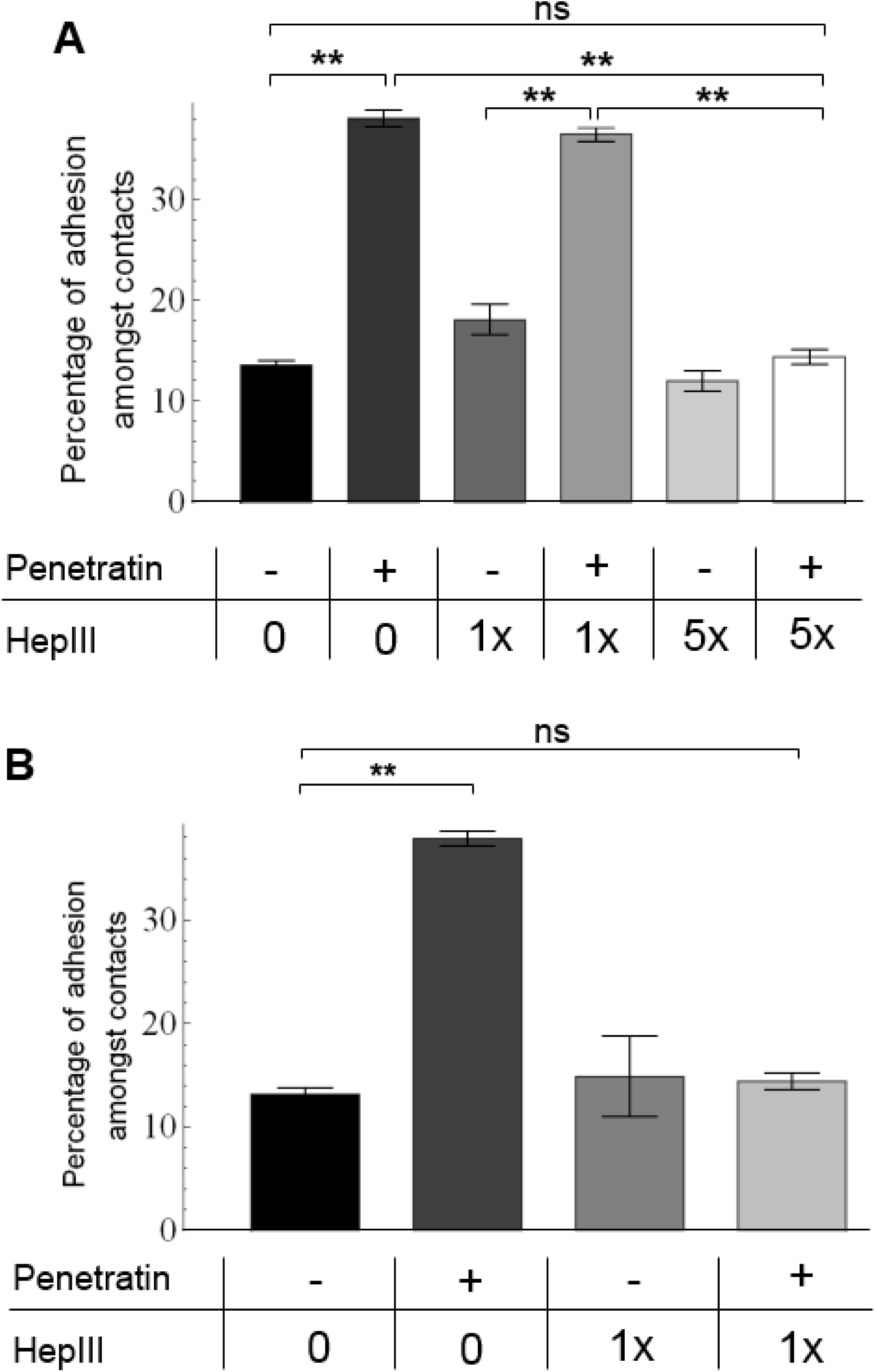
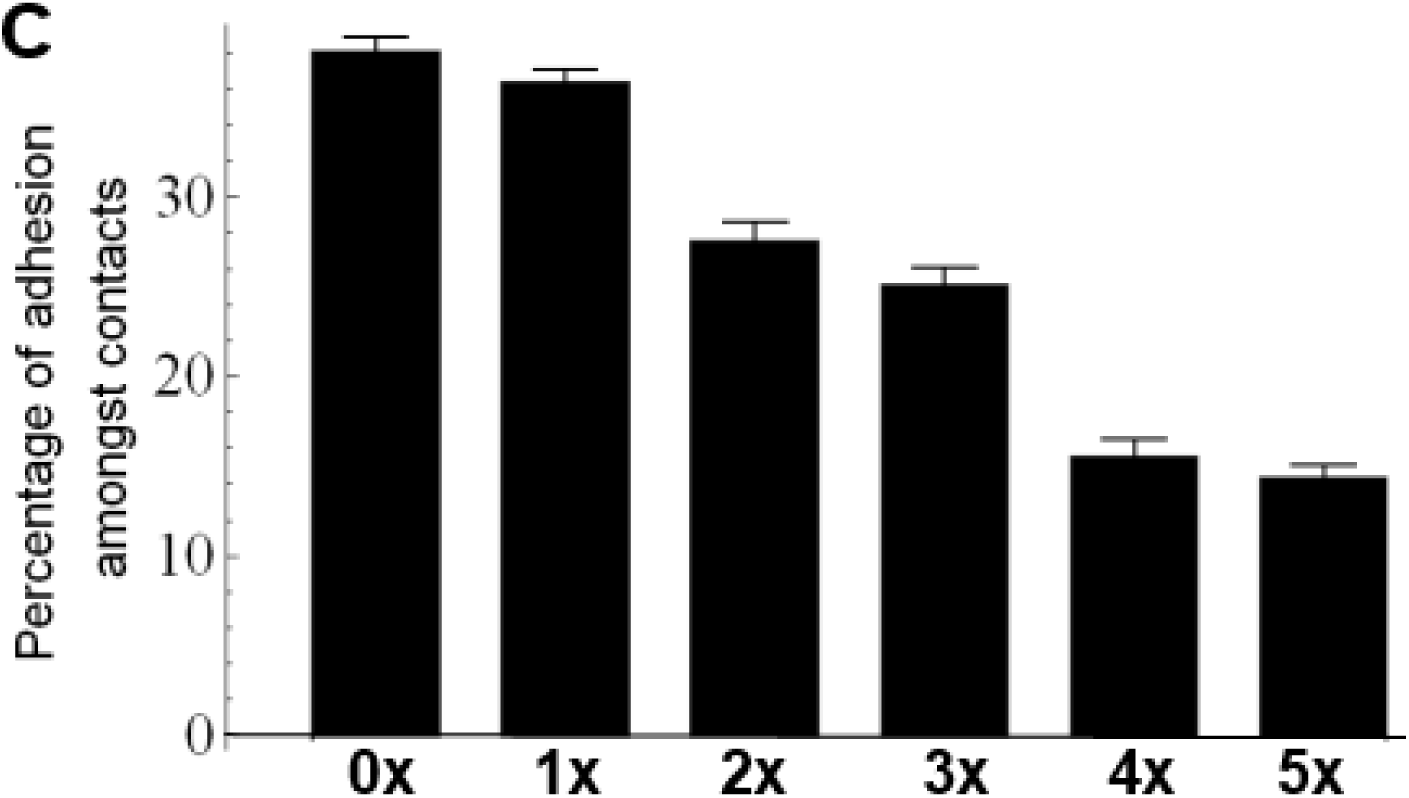
Heparan sulfates are responsible for the adhesions of Penetratin on wild type and 745 CHO cells. A: WT CHO cells B: GAG deficient CHO cells The frequency of specific adhesion is the same for wild type and GAG deficient CHO cells. These adhesions are due to heparan sulfates because they can be abolished by 1x heparinase III treatment of GAG deficient cells and 5x heparinase III treatment of wild type CHO cells. C: Dose-response effect of the heparinase III treatment of wild type CHO cells. From 1x to 5x heparinase III treatment the specific adhesion of Penetratin is gradually abolished.

Then we tested whether the specific interaction observed between Penetratin and wild-type CHO cells was also due to HSs. The same heparinase III treatment was applied on wild type CHO cells. This treatment lowered significantly (p<0.001) the specific interaction frequency to 18.4% (as compared with 24.5% in absence of heparinase III). The fact that heparinase III treatment on wild type CHO cell did not abolish completely the interactions with Penetratins can be explained by two hypotheses: (i) HSs are not the only partner of Penetratin on wild type CHO cells in contrast with GAG deficient CHO cells (ii) HSs being denser on wild type CHO cells than GAG deficient CHO cells a stronger heparinase III treatment is required to fully remove them.

To test these hypotheses, we submitted the wild type CHO cells to more stringent heparinase III treatments increasing the concentration of enzyme by a factor 2 to 5. The results are presented on figure 3B. We observed that a 5x heparinase III treatment was sufficient to fully abolish the specific interactions. From this observation we concluded that HSs are the main partner of the Penetratin-membrane interaction both on wild type CHO cells and GAG deficient CHO cells. The energy of adhesion we found is *E*_*a*_ =18 to 22*k*_*B*_*T*. In term of dissociation constant it corresponds to *K*_*d*_ = 0.3 TO 15 *nM*. This value is an approximation because the value of *E*_*a*_ is approximate and because molecules on beads are not free in solution. Nevertheless it is a comparable order of magnitude as the one found in calorimetry or PWR for Penetratin and heparan sulfate (Alves et al., 2011; Bechara et al., 2013).

### Sialic acids hide an unknown Penetratin partners

Another cell compound supposed to interact with the CPPs is the sialic acids (SAs). We disposed of another CHO mutant which is lacking SAs, the CHO Lec2 cells (SA deficient cells). The SA deficient cells did not exhibit a difference in terms of non specific interaction. They did interact a lot more with the Penetratin, since the frequency of specific adhesions between Penetratin coated beads and SA deficient cells reached 59% (figure 4). In order to determine if these interactions were due to an increase in HS accessibility we performed a heparinase III treatment on these cells. Upon 5x treatment, the amount of specific adhesion decreased by 37%, a close decrease to the rate of HS dependent specific adhesions observed on the wild type cells. However there were still 32% of unexplained specific interactions. We concluded that the absence of SA revealed other partners for the Penetratin which are not HSs.

**Figure 4:**
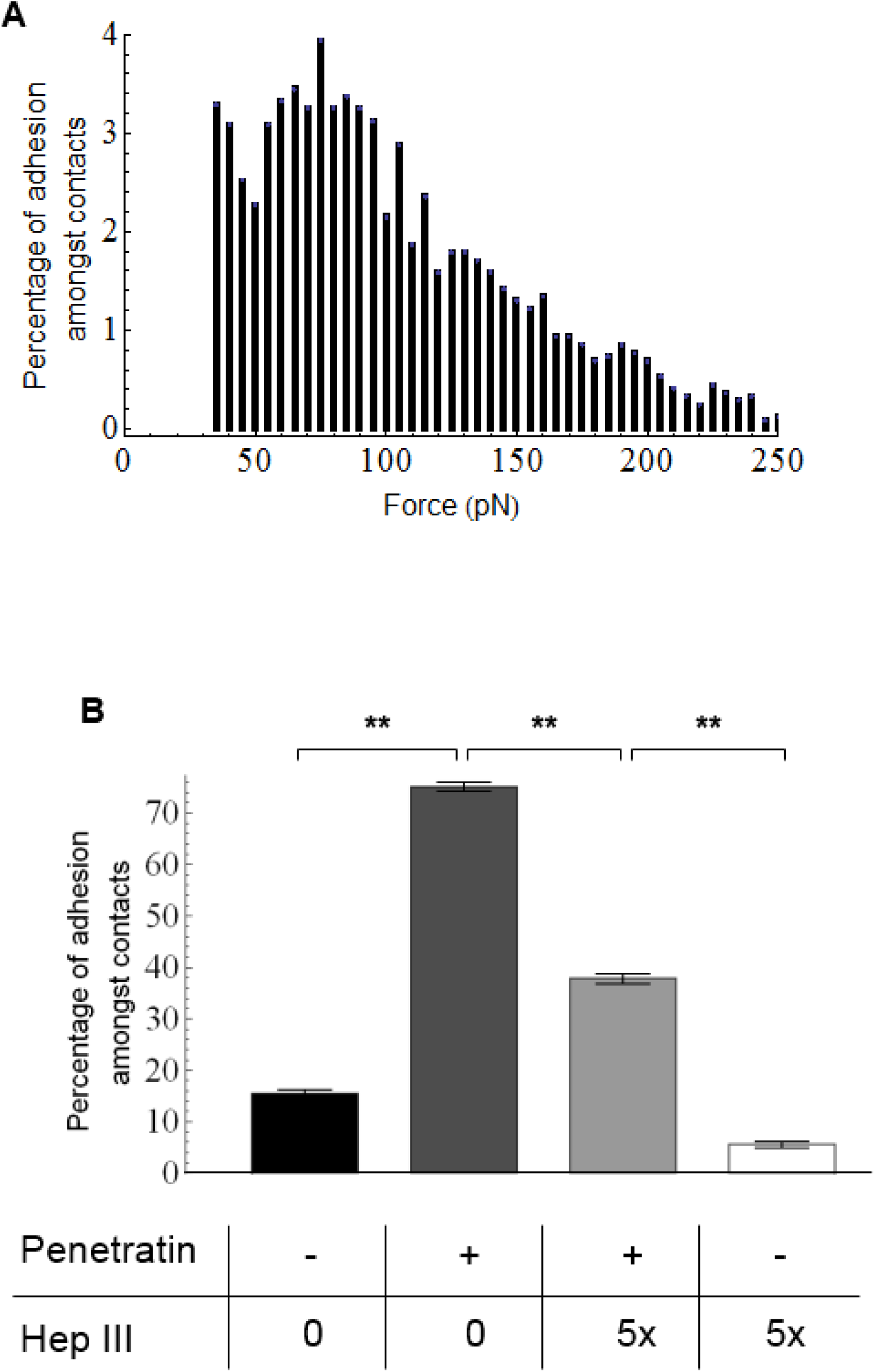
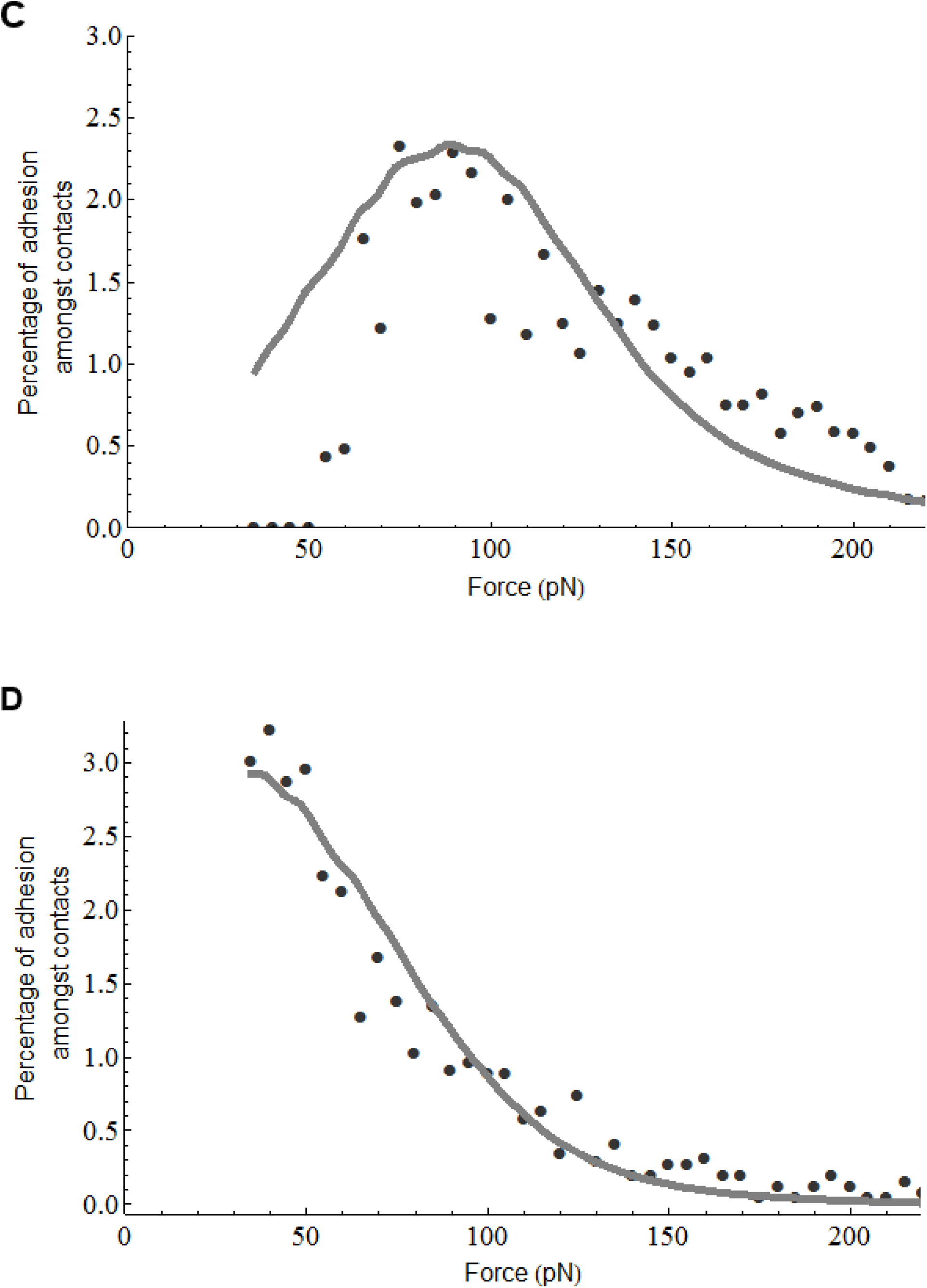
The removal of sialic acids reveals other adhesion partners different from heparan sulfates. A: Distribution of adhesion forces between Penetratin coated beads and SA deficient cells. SA deficient CHO cells exhibit a higher proportion of high adhesion forces that wild type CHO cells B: Adhesion rates on SA deficient cells. Penetratin adhered more frequently on SA deficient CHO cells than wild type CHO cells. These adhesions are only partly due to heparan sulfates showing the occurrence of other partners hidden by sialic acids. C: Comparison between the measured HS dependent force distribution on SA deficient CHO cells (black dots) and a simulated simple well model (gray line). The parameters of the best fit model are : *E*_*am*_ = 22.6 *k*_*B*_*T, σ*_*a*_ = 1.5 *k*_*B*_*T, v*_0_ = 10^10^ *s*^−1^, *p*1 = 0.36, *p*2 = 0.14 and *x*_*w*_ = 2.2 Å. D: Comparison between the measured other partner dependent force distribution on SA deficient CHO cells (black dots) and a simulated simple well model (gray line) The parameters of the best fit model are : *E*_*am*_ = 20.1 *k*_*B*_*T, σ*_*a*_ = 1.6 *k*_*B*_*T, v*_0_ = 10^10^ *s*^−1^, *p*1 = 0.36, *p*2 = 0.14 and *x*_*w*_ = 2.2 Å.

We analyzed the force distributions of the HS dependent component (obtained by substraction of the Penetratin coated beads on SA deficient cells treated with HepIII force distribution to the Penetratin coated beads on SA deficient cells without treatment) (figure 4C). This force distribution is slightly different from the HS dependent force distribution on wild type CHO cells (figure 2C). It may indicate a modification of the interactions of the Penetratin with HS in absence of SA, possibly due to a different organization of the GAGs on SA deficient cells. By using the single well modelization of the energy landscape we deduced that the energy of adhesion is 21 to 24 *k*_*B*_*T*. This is slightly higher than the HS dependant adhesion energy on WT CHO cell. It thus seems that the removal of SAs strengthen the adhesion of the Penetratin on HSs.

We also analyzed the unknown partner force distribution curve obtained by subtraction of the non covered beads on SA deficient cells treated with HepIII force distribution to the Penetratin on SA deficient cells treated with HepIII force distribution (figure 4D). We found an adhesion energy of 19 to 22 *k*_*B*_*T* for this other partner.

### A specific binding to HSs in their cellular environment rather than to any negative charges

To study the role of the electrostatic interactions between the charges of Penetratin and the charges of its cell partners, we tried to mimic these interaction using synthetic models. The aim is to determine whether charge of the partner is enough to lead to adhesion or if the nature of the chemical group bearing the charge and its environment matter.

We first tried vesicles 100% negatively charged. We made 100% DOPS giant multilamellar vesicles and tested their interaction with Penetratin coated beads and non coated beads. Despite the high charge density of these vesicles, we recorded scarce specific adhesions between Penetratin and these vesicles with a rate of 1.8% (figure 5). This suggested that charge density is not enough to provoke an adhesion.

**Figure 5:**
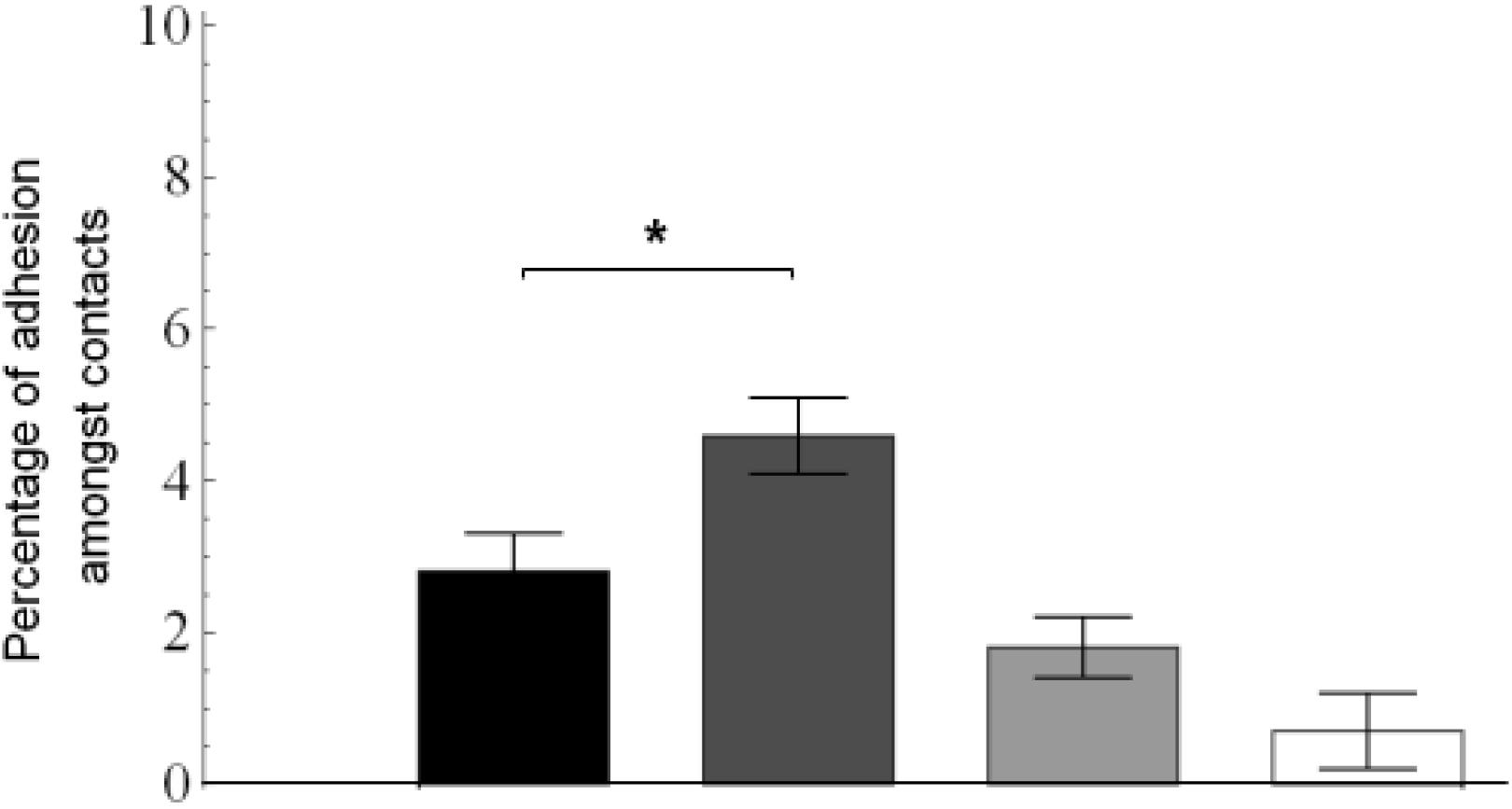
Scarce adhesion of Penetratin is observed on vesicles bearing negative charges on PS head or sulfated sugars. Black: 100% DOPS giant vesicles, non coated beads; Dark gray: 100%DOPS giant vesicles, Penetratin coated beads; Ligth gray: DOPC: sulfatides glycosphingolipids 4:1 giant vesicles, non coated beads; White: DOPC: sulfatides glycosphingolipids 4:1 giant vesicles, Penetratin coated beads The frequence of specific adhesion between giant MLV and Penetratin coated beads is below 5% for 100% DOPS and DOPC: sulfatides glycosphingolipids 4:1 vesicles.

We also tried vesicles made of 80% of DOPC lipids and 20% of lipids bearing sulfated sugars (sulfatides glycosphingolipids) in order to mimic the negative charges of the GAGs. With these vesicles, we almost recorded no adhesions between Penetratin coated beads and the vesicles (figure 5). The sulfated sugars of HSs on GAG born by cells adhere to the Penetratin but not the sulfate groups born by these lipids. This indicates a specificity of the Penetratin for the sulfated sugars born by GAG on cells and points a role for the environment of the negatively charged chemical group for the adhesion.

## CONCLUSION

We have applied the BFP technique to quantitatively study the partners of Penetratin during the first second of its arrival on a plasma membrane. Heparan sulfates are pointed as the main partner and the dynamics of sialic acids must be taken into account because they may reveal other strongly interacting partners. We have also evidenced that charge is not enough to be a strong Penetratin partner and that the chemical group bearing the charge is important.

## BIBLIOGRAPHY

Alves, I.D., Bechara, C., Walrant, A., Zaltsman, Y., Jiao, C.-Y., and Sagan, S. (2011). Relationships between Membrane Binding, Affinity and Cell Internalization Efficacy of a Cell-Penetrating Peptide: Penetratin as a Case Study. PLoS ONE 6.

Bechara, C., and Sagan, S. (2013). Cell-penetrating peptides: 20 years later, where do we stand? FEBS Lett. 587, 1693–1702.

Bechara, C., Pallerla, M., Zaltsman, Y., Burlina, F., Alves, I.D., Lequin, O., and Sagan, S. (2013). Tryptophan within basic peptide sequences triggers glycosaminoglycan-dependent endocytosis. FASEB J. Off. Publ. Fed. Am. Soc. Exp. Biol. 27, 738–749.

Burlina, F., Sagan, S., Bolbach, G., and Chassaing, G. (2005). Quantification of the cellular uptake of cell-penetrating peptides by MALDI-TOF mass spectrometry. Angew. Chem. Int. Ed Engl. 44, 4244–4247.

Burlina, F., Sagan, S., Bolbach, G., and Chassaing, G. (2006). A direct approach to quantification of the cellular uptake of cell-penetrating peptides using MALDI-TOF mass spectrometry. Nat. Protoc. 1, 200–205.

Console, S., Marty, C., García-Echeverría, C., Schwendener, R., and Ballmer-Hofer, K. (2003). Antennapedia and HIV transactivator of transcription (TAT) “protein transduction domains” promote endocytosis of high molecular weight cargo upon binding to cell surface glycosaminoglycans. J. Biol. Chem. 278, 35109–35114.

Derossi, D., Joliot, A.H., Chassaing, G., and Prochiantz, A. (1994). The third helix of the Antennapedia homeodomain translocates through biological membranes. J. Biol. Chem. 269, 10444–10450.

Dupont, E., Prochiantz, A., and Joliot, A. (2015). Penetratin Story: An Overview. Methods Mol. Biol. Clifton NJ 1324, 29–37.

Esko, J.D., Stewart, T.E., and Taylor, W.H. (1985). Animal cell mutants defective in glycosaminoglycan biosynthesis. Proc. Natl. Acad. Sci. U. S. A. 82, 3197–3201.

Evans, E., Ritchie, K., and Merkel, R. (1995). Sensitive force technique to probe molecular adhesion and structural linkages at biological interfaces. Biophys. J. 68, 2580–2587.

Evans, E., Leung, A., Heinrich, V., and Zhu, C. (2004). Mechanical switching and coupling between two dissociation pathways in a P-selectin adhesion bond. Proc. Natl. Acad. Sci. U. S. A. 101, 11281–11286.

Illien, F., Rodriguez, N., Amoura, M., Joliot, A., Pallerla, M., Cribier, S., Burlina, F., and Sagan, S. (2016). Quantitative fluorescence spectroscopy and flow cytometry analyses of cell-penetrating peptides internalization pathways: optimization, pitfalls, comparison with mass spectrometry quantification. Sci. Rep. 6, 36938.

Jégou, A., Pincet, F., Perez, E., Wolf, J.P., Ziyyat, A., and Gourier, C. (2008). Mapping mouse gamete interaction forces reveal several oocyte membrane regions with different mechanical and adhesive properties. Langmuir ACS J. Surf. Colloids 24, 1451–1458.

Jiao, C.-Y., Delaroche, D., Burlina, F., Alves, I.D., Chassaing, G., and Sagan, S. (2009). Translocation and endocytosis for cell-penetrating peptide internalization. J. Biol. Chem. 284, 33957–33965.

Kurrikoff, K., Veiman, K.-L., and Langel, Ü. (2015). CPP-Based Delivery System for In Vivo Gene Delivery. Methods Mol. Biol. Clifton NJ 1324, 339–347.

Kurrikoff, K., Gestin, M., and Langel, Ü. (2016). Recent in vivo advances in cell-penetrating peptide-assisted drug delivery. Expert Opin. Drug Deliv. 13, 373–387.

Merkel, R., Nassoy, P., Leung, A., Ritchie, K., and Evans, E. (1999). Energy landscapes of receptor-ligand bonds explored with dynamic force spectroscopy. Nature 397, 50–53.

Pincet, F., and Husson, J. (2005). The Solution to the Streptavidin-Biotin Paradox: The Influence of History on the Strength of Single Molecular Bonds. Biophys. J. 89, 4374–4381.

Rothbard, J.B., Jessop, T.C., Lewis, R.S., Murray, B.A., and Wender, P.A. (2004). Role of membrane potential and hydrogen bonding in the mechanism of translocation of guanidinium-rich peptides into cells. J. Am. Chem. Soc. 126, 9506–9507.

Ziegler, A., and Seelig, J. (2008). Binding and Clustering of Glycosaminoglycans: A Common Property of Mono- and Multivalent Cell-Penetrating Compounds. Biophys. J. 94, 2142–2149.

